# Human REXO4 is Required for Cell Cycle Progression

**DOI:** 10.1101/2025.01.08.631954

**Authors:** Kevin M. Clutario, Mai Abdusamad, Ivan Ramirez, Kayla J. Rich, Ankur A. Gholkar, Julian Zaragoza, Jorge Z. Torres

## Abstract

Human REXO4 is a poorly characterized exonuclease that is overexpressed in human cancers. To better understand the function of REXO4 and its relationship to cellular proliferation, we have undertaken multidisciplinary approaches to characterize its cell cycle phase-dependent subcellular localization and the cis determinants required for this localization, its importance to cell cycle progression and cell viability, its protein-protein association network, and its activity. We show that the localization of REXO4 to the nucleolus in interphase depends on an N-terminal nucleolar localization sequence and that its localization to the perichromosomal layer of mitotic chromosomes is dependent on Ki67. Depletion of REXO4 led to a G1/S cell cycle arrest, and reduced cell viability. REXO4 associated with ribosome components and other proteins involved in rRNA metabolism. We propose a model where REXO4 is important for proper rRNA processing, which is required for ribosome biogenesis, cell cycle progression, and proliferation.

**SIGNIFICANCE STATEMENT:**

- REXO4 is a putative RNA exonuclease with limited characterization. The authors used in silico, cell, and molecular biology approaches to characterize its localization, associations, regulation, and function.
- They found that during interphase, REXO4 localizes to the nucleolus through an N-terminal nucleolar localization sequence. Whereas during mitosis, REXO4 localized to the perichromosomal layer in a Ki67-dependent manner. REXO4 was required for proper cell cycle progression, and viability.
- These results indicated that REXO4 is important for regulating cell proliferation.

## INTRODUCTION

Ribosomes are macromolecular complexes comprised of RNAs and proteins that translate messenger RNAs (mRNAs) into amino acid polypeptides by providing a catalytic site for peptide bond formation (Dorner *et al*., 2023). The biogenesis of ribosomes is a complex process requiring not only the expression, modification, and regulation of about 80 ribosomal proteins, but also the expression, modification, and processing of four ribosomal RNAs (rRNAs) (Dorner *et al*., 2023). In humans, RNA polymerase I (POLR1A) is responsible for transcribing a 47S pre-rRNA precursor from hundreds of ribosomal DNA (rDNA) repeats located on the five acrocentric chromosomes within the nucleolus (Grummt, 2003). The 47S pre-rRNA precursor is processed into three of the four rRNAs (28S, 5.8S, and 18S) incorporated into ribosomes (Aubert *et al*., 2018). Whereas RNA polymerase III (POLR3A) is responsible for transcribing the pre-5S rRNA precursor from chromosome 1, which is processed into the 5S rRNA (Watt *et al*., 2023). The processing and maturation of the pre-rRNA into mature rRNAs relies on a series of processing steps carried out by multiple endo- and exonucleases (Henras *et al*., 2015). While much is known about the processing and maturation steps of pre-rRNA into mature rRNAs, the full complement of ribonucleolytic enzymes and how they are regulated to carry out this complex process with high fidelity is not fully understood.

RNA metabolism relies heavily on exoribonucleases, which are subdivided into the RNR, DEDD, RBN, PDX, RRP4, and 5PX superfamilies (Zuo and Deutscher, 2001). Members of the DEDD superfamily are characterized by four conserved acidic residues (DEDD) found within the three exonuclease motifs (Exo I-III) of their exonuclease domain (Zuo and Deutscher, 2001). Further, the DEDDh subfamily members, within the DEDD superfamily, also have a fifth conserved residue (histidine, H) in the Exo III motif that distinguishes them from DEDDy subfamily members that have a conserved tyrosine (Y) at this site (Zerbino *et al*., 2018). The Rex/Rexo group of proteins are RNA exonucleases within the DEDDh subfamily that have been shown to have key roles in RNA processing. For example, in *Saccharomyces cerevisiae*, Rex1p, Rex2p, and Rex3p are involved in the processing of non-coding small nuclear RNA (snRNA) and rRNA (van Hoof *et al*., 2000). Rex2p and Rex3p have also been implicated in processing coding mature RNA (mRNA) (Hodko *et al*., 2016). In *Drosophila melanogaster*, Rexo5 was shown to be important for processing of the 28S and 5S rRNAs and small nucleolar RNA (snoRNA) (Gerstberger *et al*., 2017).

Another Rex/Rexo group member was initially described in *Xenopus laevis* as *<u>X</u>enopus* <u>p</u>revents <u>m</u>itotic <u>c</u>atastrophe 2 (XPMC2), a gene that encoded a nuclear protein that could rescue the mitotic catastrophe observed in Wee1 kinase mutants in *Schizosaccharomyces pombe (Su and Maller, 1995)*. The human homolog of XPMC2 was then cloned and named *<u>X</u>enopus* <u>p</u>revents <u>m</u>itotic <u>c</u>atastrophe 2 homolog (XPMC2H) (Kwiatkowska *et al*., 1997) and was later identified as a factor that modulated quinone reductase gene transcriptional activity and named <u>h</u>uman homolog of *Xenopus* gene which <u>p</u>revents <u>m</u>itotic <u>c</u>atastrophe (hPMC2) (Montano *et al*., 2000) and subsequently renamed RNA Exonuclease 4 (REXO4) based on sequence homology to the Rex/Rexo exoribonucleases. Although human REXO4 was shown to possess 3′ to 5′ non-processive exonuclease activity that degrades both single-stranded and double-stranded DNA (Krishnamurthy *et al*., 2011), it has remained poorly characterized regarding its subcellular localization, regulation, and function. Moreover, the REXO4 homolog in *S. cerevisiae*, Rex4p, was genetically linked to the processing of the first internal transcribed spacer ITS1 within the 47S pre-rRNA and regulation of the relative levels of 5.8S rRNA short form (5.8S_S_) and long form (5.8S_L_) (Faber *et al*., 2004). However, human REXO4’s role in RNA processing has not been explored. Recent reports indicate that REXO4 expression levels are a biomarker for hepatocellular carcinoma disease progression and outcome (Chen *et al*., 2021; Ruan *et al*., 2021). Therefore, further characterization of REXO4’s function and regulation are important for understanding cell cycle control and proliferation.

## RESULTS

### *In silico* analysis of REXO4

The human REXO4 gene is located on the 9q34.2 region of chromosome nine and encodes for a 422 amino acid protein with a distinctive exonuclease domain near its C-terminus (Figure 1A and Supplemental Figure S1). The REXO4 protein belongs to the Rex sub-family of DEDDh exoribonucleases within the DEDD superfamily of exoribonucleases and contains the four conserved acidic residues (DEDD) found within the Exo I-III motifs of the exonuclease domain and a fifth conserved residue (H) in the Exo III motif (Supplemental Figure S1) (Zuo and Deutscher, 2001). REXO4 is conserved across eukaryotes and shares high amino acid sequence identity with homologs found in model organisms like *M. musculus* Rexo4 (70%), *X. laevis* rexo4 (55%), *D. melanogaster* CC6833 (52%), and *S. cerevisiae* REX4 (48%) (Figure 1A and Supplemental Figure S1). In humans, there are four predicted REXO4 mRNA isoforms: isoform 1 is the longest isoform that encodes a 420aa REXO4 protein; isoform 2 lacks two coding exons that results in a frameshift and is shorter than isoform 1, encodes for a 250aa REXO4 protein; isoform 3 uses an alternative splice site near the 5’ end and initiates translation at a downstream start codon resulting in a shorter 285aa REXO4 protein that lacks a portion of the N-terminus; isoform 4 is similar to isoform 3 and initiates translation at a downstream start codon resulting in a shorter 329aa REXO4 protein that lacks a portion of the N-terminus (Supplemental Figure S2). Interestingly, isoform 2 lacks the Exo I motif of the exonuclease domain and is not predicted to have exonuclease catalytic activity (Supplemental Figure S2). AlphaFold (Varadi *et al*., 2022) structural analysis of isoform 1 REXO4 showed that its N-terminus (1-243aa) contained long stretches of disorder, which was followed by a structured globular exonuclease domain (244-394aa), and a short largely unstructured C-terminus (395-422aa) (Figure 1B). Analysis of REXO4 expression across 54 non-diseased tissue types using the Adult Genotype Tissue Expression (GTEx) Portal (Consortium, 2020) showed that REXO4 was expressed in all tissues tested (Supplemental Figure S3A). Similarly, analysis of REXO4 expression through the Human Protein Atlas showed that it was expressed across 1206 human cell lines, including 1132 cancer cell lines, and clustered with genes involved in basic cellular processes (Supplemental Figure S3B) (Uhlen *et al*., 2015; Jin *et al*., 2023). Two recent reports determined that REXO4 is overexpressed in hepatocellular carcinoma (HCC) and could be used as a prognostic marker of the disease (Chen *et al*., 2021; Ruan *et al*., 2021). Further, another study found that the downregulation of REXO4 (aka hPMC2) sensitized MDA-MB-231 and MDA-MB-468 breast cancer cells to alkylating agents (Krishnamurthy *et al*., 2015). Therefore, we asked if REXO4 expression levels were widely misregulated in cancer by analyzing REXO4 expression levels across a diverse set of cancers using the Gene Expression Profiling and Interactive Analysis (GEPIA) web server (Tang *et al*., 2017). This analysis showed that REXO4 mRNA levels were elevated in most cancers analyzed compared to matched normal samples, but significant in only thymoma and lymphoid neoplasm diffuse large B-cell lymphoma (Supplemental Figure 3C and Supplemental Table S1). Together, these analyses showed that REXO4 is a highly conserved protein among eukaryotes, is expressed in all tissues and cell lines tested, and that misregulation of REXO4 levels is found in only a few types of cancers.

**Figure 1.**
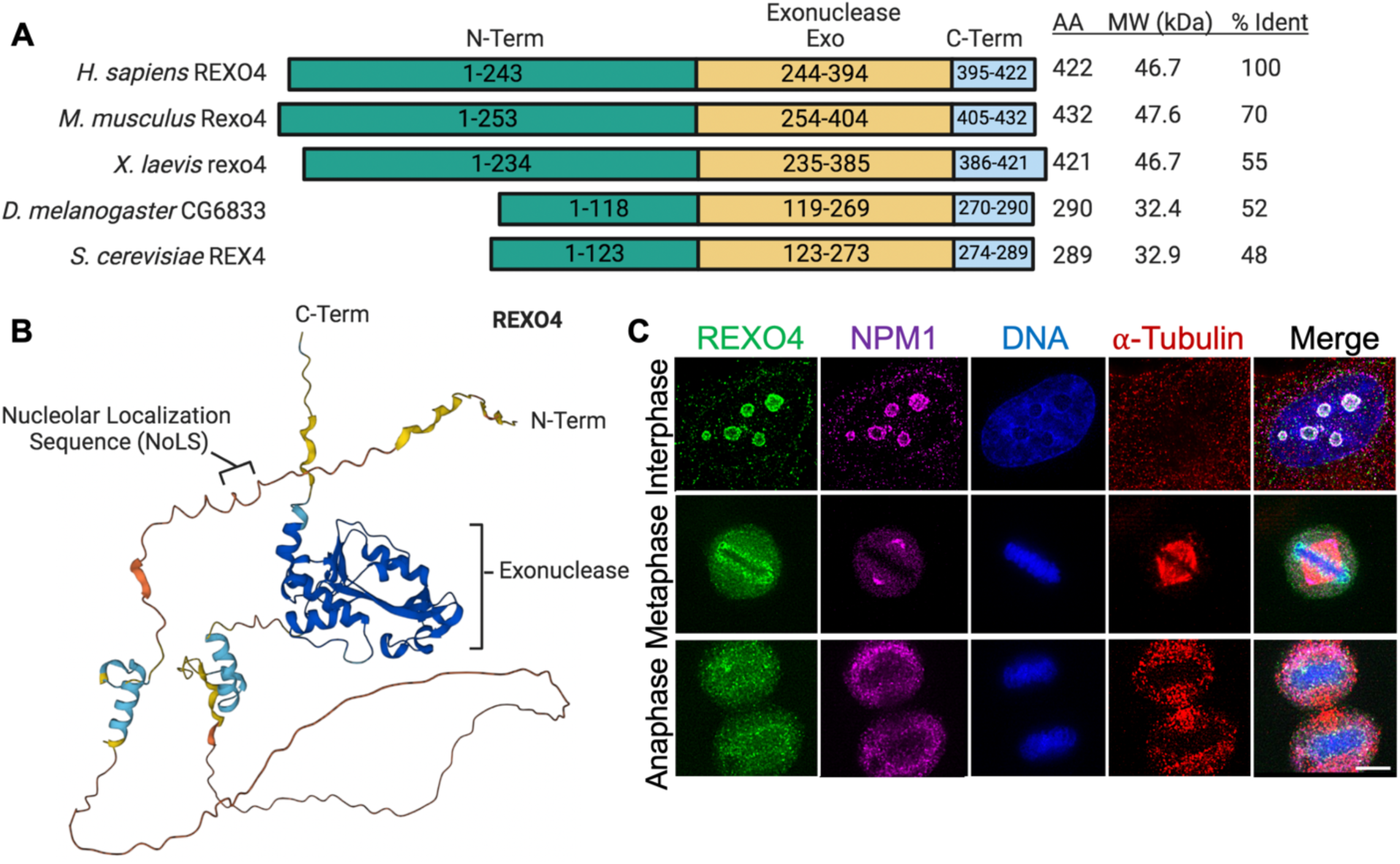
*In silico* and in cell analysis of REXO4. (A) Schematic of the protein domain architecture for human REXO4 and its eukaryotic orthologs; *H. sapiens* REXO4: UniProt ID Q9GZR2, *M. musculus* Rexo4: UniProt ID Q6PAQ4, *X. laevis rexo4*: UniProt ID Q91560, *D. melanogaster* CG6833: UniProt ID Q9VUC3, *S. cerevisiae* REX4: UniProt ID Q08237. Number of amino acids (AA), molecular weight in kilodaltons (MW kDa) and the percent amino acid identity (% Ident) with reference to human REXO4 are indicated on the right. The amino acids within the N-terminus (N-term), Exonuclease (Exo), and C-terminus (C-term) domains are as indicated. (B) Schematic of the protein domain architecture for the four predicted human REXO4 isoforms. Note that isoform 2 encodes a protein that is missing part of the Exo domain and contains a stretch of amino acids (76-132) not found in other isoforms (light yellow box). (C) Analysis of endogenous REXO4 subcellular localization throughout the cell cycle. HeLa cells were fixed with paraformaldehyde; co-stained for DNA (Hoechst 33342), tubulin (anti-α-tubulin), REXO4 (anit-REXO4), and NPM1 (anit-NPM1); and imaged by immunofluorescence microscopy. Bar indicates 5μm.

### REXO4 localizes to the nucleolus in interphase and mitotic chromosomes during cell division

To better understand the function of REXO4, we first sought to determine its cell cycle subcellular localization. HeLa cells were fixed and stained with Hoechst 3442 (DNA), anti-Tubulin antibodies, and anti-REXO4 antibodies. Cells were then imaged by immunofluorescence (IF) microscopy. This analysis showed that REXO4 likely localized to the nucleolus during interphase and to the periphery of condensed mitotic chromosomes during cell division (Figure 1C). REXO4’s nucleolar localization pattern appeared similar to the nucleolar marker NPM1 (Nucleophosmin) (Ochs *et al*., 1985). Indeed, co-staining for REXO4 and NPM1 showed that their localizations overlapped at the nucleolus in interphase (Figure 1C). To better visualize the REXO4 localization during cell division, cells were arrested with the Kif11/Eg5 kinesin inhibitor monastrol, which arrests cells with a monopolar spindle in early prometaphase with condensed chromosomes in a rosette-like pattern (Mayer *et al*., 1999). Cells were then stained for REXO4, DNA, Tubulin, and NPM1. Consistently, REXO4 localized to the periphery of individual condensed early mitotic chromosomes (Figure 2A). The perichromosomal layer (PCL) is comprised of numerous proteins and rRNAs that coat mitotic chromosomes and it is important for chromosome compaction in mitosis and proper cell division (Van Hooser *et al*., 2005). Ki67 is a nucleolar protein that localizes to the periphery of chromosomes in early mitosis and is important for building and anchoring the PCL (Cuylen *et al*., 2016; Hayashi *et al*., 2017). To determine if REXO4 was a component of the PCL, we analyzed the localization of endogenous REXO4 and overexpressed GFP-REXO4 in a verified HeLa Ki67 knockout cell line (Figure 2B-D) (Cuylen *et al*., 2016). Interestingly, both endogenous REXO4 and overexpressed GFP-REXO4 remained localized to the nucleolus during interphase (Figure 2C). However, both endogenous REXO4 and overexpressed GFP-REXO4 were no longer localized throughout mitotic chromosomes and instead were aggregated and formed a cap at one end of the metaphase plate (Figure 2D). These results indicated that REXO4 had a dynamic cell cycle phase subcellular distribution, localizing to the nucleolus in interphase and to mitotic chromosomes during mitosis. Additionally, our results showed that Ki67 was required for proper REXO4 localization to mitotic chromosomes and that it was a component of the PCL.

**Figure 2.**
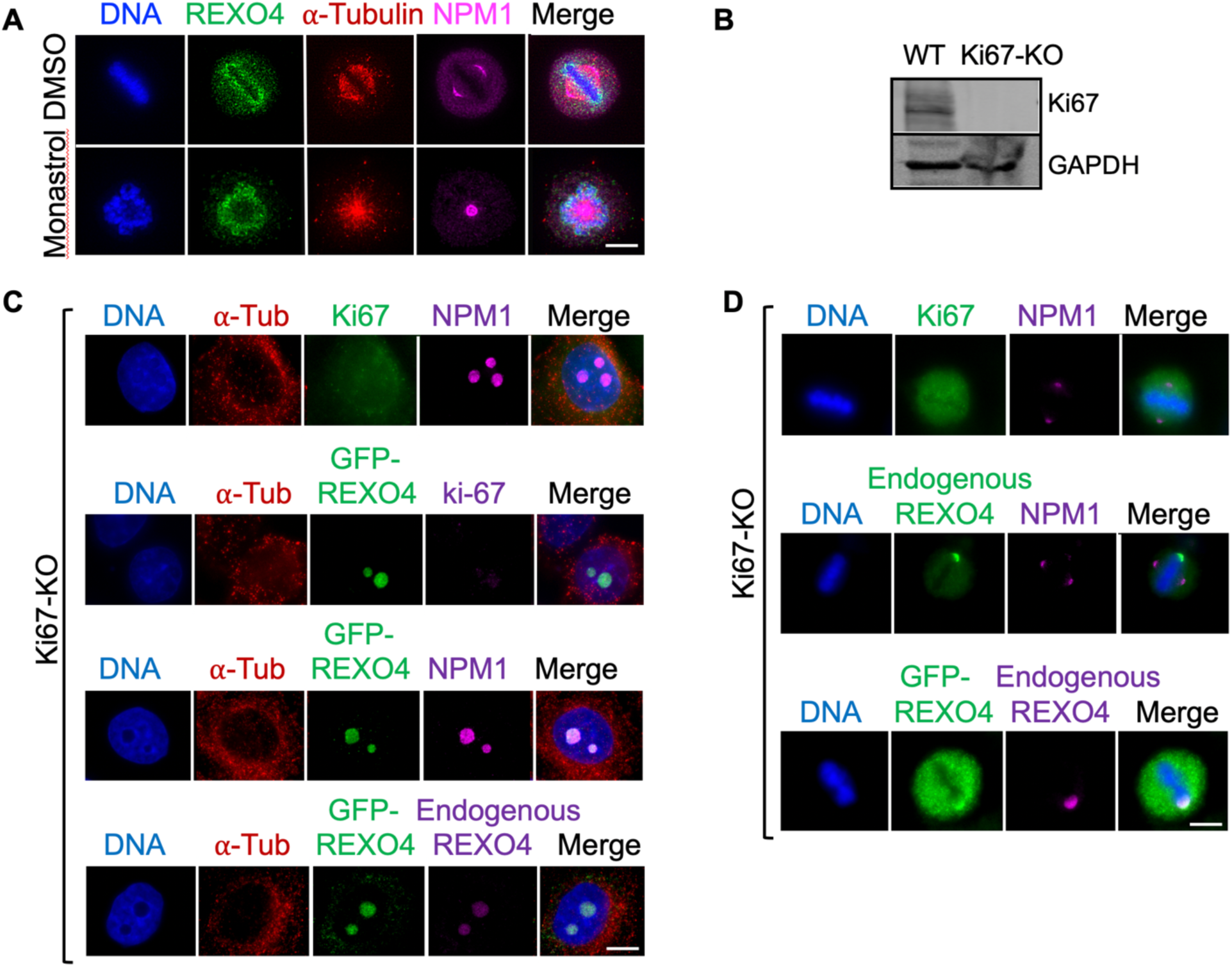
REXO4 localizes to the nucleolus in interphase cells in a Ki67 independent manner and to mitotic chromosomes during cell division in a Ki67 dependent manner. (A) Analysis of endogenous REXO4 subcellular localization in cells treated with +/- 50 μM monastrol for 16 hours. Cells were fixed with paraformaldehyde; co-stained for DNA (Hoechst 33342), tubulin (anti-α-tubulin), REXO4 (anit-REXO4), and NPM1 (anit-NPM1); and imaged by immunofluorescence microscopy. Bar indicates 5μm. (B) Immunoblot analysis of protein extracts from HeLa WT or Ki67 knock out (Ki67-KO) cells using anti-GAPDH and anti-Ki67 antibodies. (C-D) Ki67-KO cells were transfected with +/- full-length GFP-REXO4, fixed, and co-stained for DNA (Hoechst 33342), tubulin (anti-α-tubulin), REXO4 (anit-REXO4), NPM1 (anit-NPM1), Ki67 (anti-Ki67), or GFP (anti-GFP to detect GFP-REXO4) as indicated. Bar indicates 5μm.

To better understand the dynamic localization of REXO4 to the nucleolus and mitotic chromosomes and the cis determinants required for this localization, we generated GFP-tagged versions of REXO4 full-length (FL) and a series of truncation mutants that included the N-terminal (NT, aa1-243) domain, the exonuclease (Exo, aa244-394) domain, and the exonuclease- C-terminal (ExoCT, aa244-422) fragment (Figure 3A). GFP-REXO4 FL, NT, Exo, and ExoCT were transiently expressed in HeLa cells, which were fixed and stained for DNA, Tubulin, GFP, and NPM1. While REXO4 FL and NT localized properly to the nucleolus during interphase, Exo and ExoCT failed to localize to the nucleolus (Figure 3B). The localization of REXO4 FL to the nucleolus was consistent with a previous proteome-wide GFP-tagging protein subcellular localization campaign that showed GFP-REXO4 localized to the nucleus and nucleolus (Simpson *et al*., 2000). Interestingly, while GFP-REXO4 FL localized properly to mitotic chromosomes, all other fragments were predominantly dispersed in mitotic cells (Supplemental Figure S4). These results indicated that the REXO4 N-terminus harbored determinants for nucleolar localization and that full-length REXO4 was necessary for proper localization to mitotic chromosomes.

**Figure 3.**
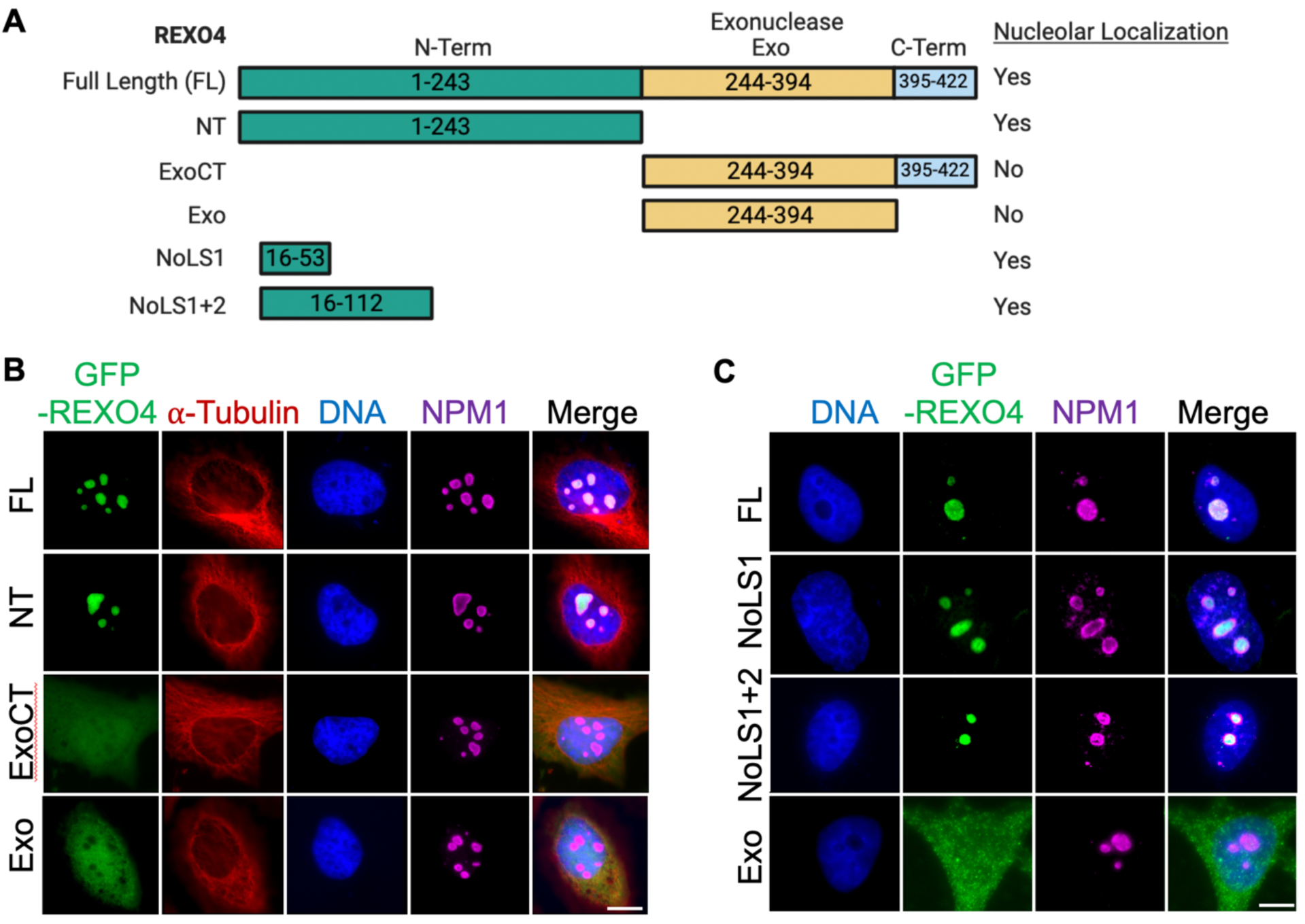
REXO4 targets to the nucleolus through an N-terminal nucleolar localization sequence. (A) Schematic of REXO4 full-length (FL), and the N-terminus (NT), Exonuclease-C-terminus (ExoCT), Exonuclease (Exo), Nucleolar localization sequence 1 (NoLS1), and Nucleolar localization sequences 1 and 2 (NoLS1+2) truncation mutants that were cloned into the mammalian expression vector pgLAP1 (N-terminal GFP-S-peptide tag) to generate inducible HeLa stable cell lines capable of expressing these tagged proteins. (B-C) HeLa cells were induced to express GFP-REXO4-FL, GFP-REXO4-NT, GFP-REXO4-ExoCT, GFP-REXO4-Exo, GFP-REXO4-NoLS1, or GFP-REXO4-NoLS1+2 for 16 hours prior to fixation. Cells were stained for DNA (Hoechst 33342) and either GFP-REXO4 (anti-GFP), tubulin (anti-α-tubulin), or NPM1 (anit-NPM1) as indicated. Bar indicates 5μm.

### REXO4 is targeted to the nucleolus through an N-terminal nucleolar localization sequence

Since the N-terminal region of REXO4 was important for its nucleolar localization, we analyzed the REXO4 protein for any putative nucleolar localization sequences (NoLSs) using the NoD (<u>n</u>ucle<u>o</u>lar localization sequence <u>d</u>etector) program (Scott *et al*., 2010; Scott *et al*., 2011). This analysis revealed a strong putative NoLS at the N-terminus of REXO4, and two weak NoLSs; one in the N-terminus and one within the Exo domain (Supplemental Figure S5). Given that the Exo and ExoCT REXO4 fragments did not localize properly to the nucleolus and that the NT fragment did (Figure 3B), we asked whether one or both of the N-terminal NoLSs were sufficient to localize REXO4 to the nucleolus. The two NoLSs were within amino acids 16-112 (NoLS1: 16-53, NoLS2: 92-112) (Supplemental Figure S5). Therefore, we generated GFP-tagged REXO4 NoLS1+2 (aa 16-112) and NoLS1 (aa16-53), and asked if they could target GFP to the nucleolus. Indeed, similar to REXO4 full-length, REXO4 NoLS1+2 and NoLS1, were sufficient to localize GFP to the nucleolus (Figure 3C). Together, these results indicated that REXO4 has a NoLS within its N-terminal domain that is important for its nucleolar targeting.

### REXO4 is important for cell cycle progression

REXO4 expression has been proposed as a prognostic marker of hepatocellular carcinoma (Chen *et al*., 2021; Ruan *et al*., 2021) and our analysis showed that high REXO4 expression levels are also found elevated in various cancers and significantly in thymoma and lymphoid neoplasm diffuse large B-cell lymphoma. (Supplemental Figure 3C). Therefore, we sought to determine the effect of depleting REXO4 on the cell cycle and cell proliferation. First, HeLa cells were treated with control non-targeting siRNA (siNT) or siRNA targeting REXO4 (siREXO4-1 and siREXO4-2) for 72 hours, and protein extracts were analyzed by immunoblot analysis. Both REXO4 siRNAs depleted REXO4 efficiently compared to the non-targeting control (Figure 4A). Next, we analyzed whether depletion of REXO4 affected cell viability using the CellTiter-Glo 2.0 cell viability luminescent assay that measures the levels of ATP, indicative of metabolically active cells. This assay showed that treatment of cells with siREXO4 led to a marked decrease in the percent cell viability at 72 hours hours post-treatment compared to siControl (Figure 4B). Next, we analyzed whether the depletion of REXO4 was affecting the cell cycle. HeLa cells were treated with siControl or siREXO4 for 72 hours and analyzed by flow cytometry. This analysis showed that siREXO4 treated cells were accumulating in G1 phase (Figure 4C). Together, these results indicated that depletion of REXO4 arrested cells in G1 phase and reduced cell viability.

**Figure 4.**
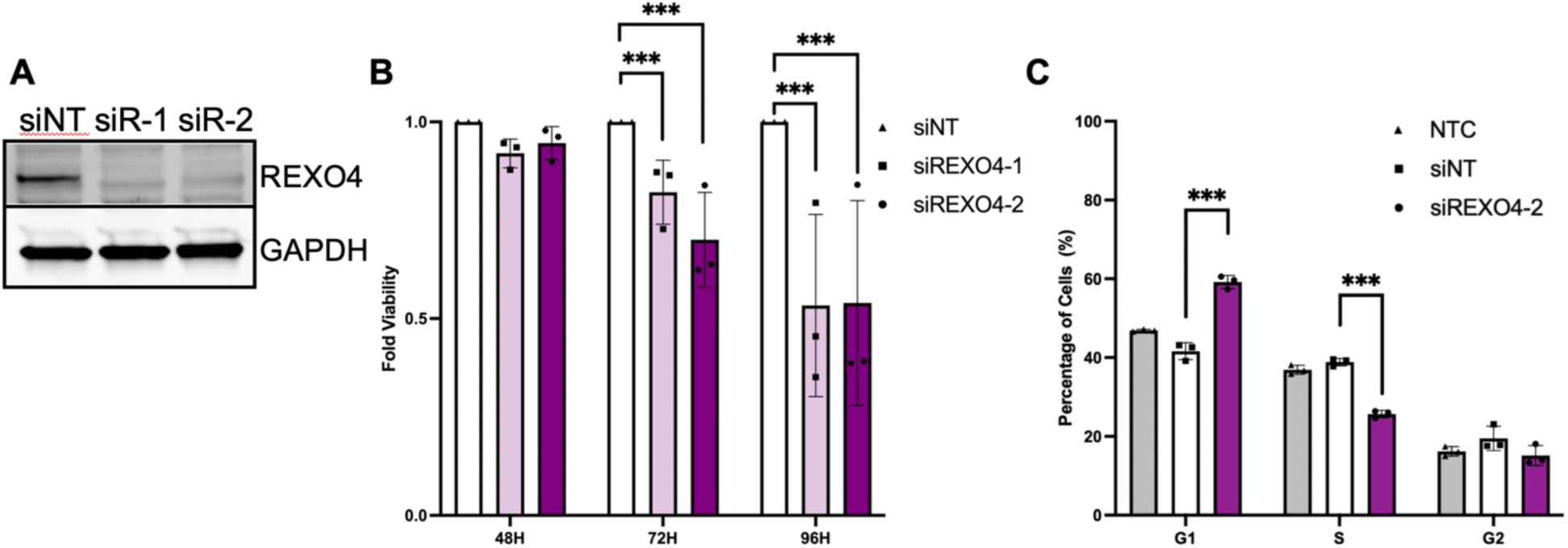
Depletion of REXO4 leads to cell cycle arrest and decreased cell viability. (A) Immunoblot analysis of protein extracts from HeLa cells treated with non-targeting control siRNA (siControl) or siRNA targeting REXO4 (siREXO4-1 and siREXO4-2) for 72 hours. (B) HeLa cells were treated with siControl or siREXO4 for 48, 72, or 96 hours and their viability was assessed with the CellTiter-Glo assay. Y-axis represents fold viability normalized to siControl within each timepoint. Data represent the average ± SD of three independent experiments with three technical triplicates each normalized to siControl. *** indicates a p value =<.0001. (C) HeLa cells were treated with siControl or siREXO4 for 72 hours and subjected to FACS cell cycle analysis. Data represent the average ± SD of three independent experiments, 1000 cells counted for each. *** indicates a p value =<.0001.

### REXO4 associates with ribosome and RNA metabolism proteins

To better understand the function of REXO4, we generated a LAP (Localization and Affinity Purification; GFP-S-protein) tagged REXO4 doxycycline (Dox) inducible HEK293T Flp-In T-REx cell line to define REXO4’s protein association network (Supplemental Figure 6A) (Torres *et al*., 2009; Bradley *et al*., 2016). Briefly, the LAP-REXO4 cell line was Dox-induced for 24 hours, cells were harvested, and protein extracts were prepared. LAP-REXO4 was tandem affinity purified, GFP first followed by S-protein, and the purified eluates were trypsin digested and analyzed by LC-MS/MS. A LAP-tag only data set was used to subtract common contaminants from the LAP-REXO4 data set to establish a list of proteins representing the REXO4 association network (Supplemental Table S2). The REXO4 association network was generated using CANVS (<u>C</u>lean <u>A</u>nalyze <u>N</u>etwork <u>V</u>isualization <u>S</u>oftware (Velasquez *et al*., 2021). This analysis showed that REXO4 was associating with ribosome and rRNA metabolism proteins (Supplemental Figure 6B). To better visualize this data, CANVS was used to analyze the REXO4 associating proteins with the CORUM (Giurgiu *et al*., 2019), Gene Ontology (Ashburner *et al*., 2000), BioGRID (Stark *et al*., 2006), and Reactome (Jassal *et al*., 2020) databases using ribosome (GO:0005840), ribosome biogenesis (GO:0042254), rRNA processing (GO:0006364), rRNA binding (GO:0019843), and rRNA processing (GO:0006364) GO terms (Figure 5A-D). Several insights were gleaned from these analyses, first REXO4 was predominantly associated with rRNA binding and rRNA processing proteins. Second, many of these proteins were known to localize to or have a function in the nucleolus. Third, these proteins were broadly involved in ribosome biogenesis and ribosome function. Together, these results indicated that REXO4 likely had a role in rRNA processing and ribosome biogenesis.

**Figure 5.**
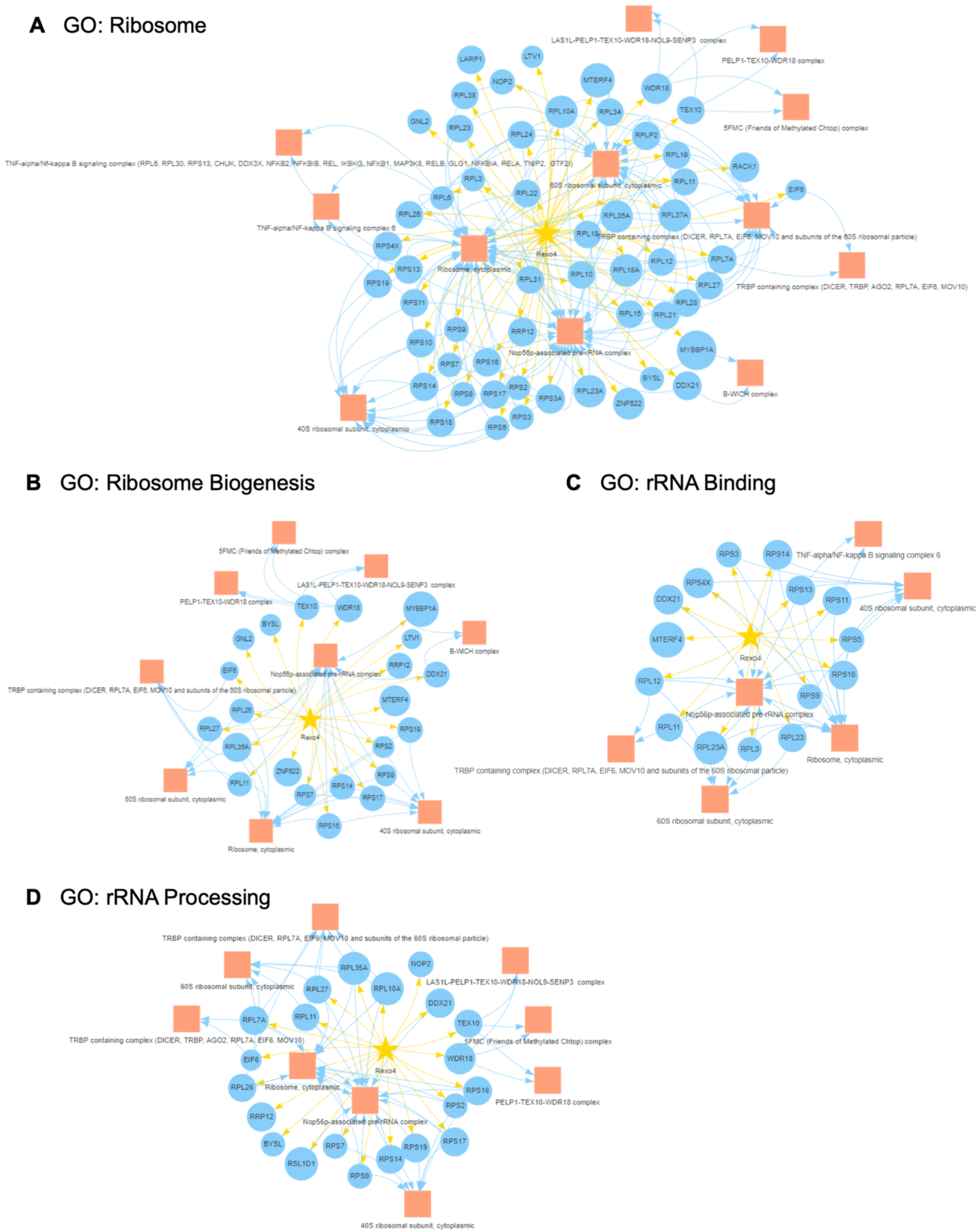
REXO4 associates with rRNA processing and ribosome biogenesis proteins. (A-D) A REXO4-LAP (GFP-S-protein) tagged HEK293T Flp-In T-REx cell line used for REXO4 tandem affinity purification and interacting proteins were identified by liquid chromatography tandem mass spectrometry (LC-MS/MS). Clean Analyze Network Visualization Software (CANVS) was then used to generate REXO4 protein-protein association networks using ribosome (GO:0005840), ribosome biogenesis (GO:0042254), rRNA processing (GO:0006364), rRNA binding (GO:0019843), and rRNA processing (GO:0006364) Gene Ontology (GO) terms. See Supplemental Table S2 for a complete list of identified proteins.

### REXO4 is dispersed throughout the cell in response to nucleolar stress

Previous studies have shown that a subset of nucleolar proteins like p62 (SQSTM1) and PICT1 are degraded in the presence of nucleolar stress (Maehama *et al*., 2014; Katagiri *et al*., 2015). This is thought to be a response to down-regulate proteins important for cellular proliferation under non-favorable conditions. Therefore, we sought to determine if REXO4 protein levels were responsive to nucleolar stress. HeLa cells were treated for 24 hours with DMSO control vehicle or 5 nM Actinomycin D (ActD), which inhibits transcription, induces nucleolar stress, and disrupts nucleolar structure. Immunoblot analysis of protein extracts from these cells showed that PICT1 protein levels were reduced in ActD-treated cells compared to the control, while REXO4 levels remained constant (Figure 6A). This was consistent with a previous study that determined that PICT1 was degraded in response to ActD treatment (Maehama *et al*., 2014). Immunofluorescence microscopy of cells stained for REXO4, DNA, and NPM1 showed that nucleolar structures were absent in ActD-treated cells and that REXO4 was dispersed throughout the cell, similar to NPM1 (Figure 6B). Together, these results showed that the REXO4 protein levels did not decrease in response to nucleolar stress, but that REXO4 was dispersed throughout the cell in the absence of defined nucleolar structures.

**Figure 6.**
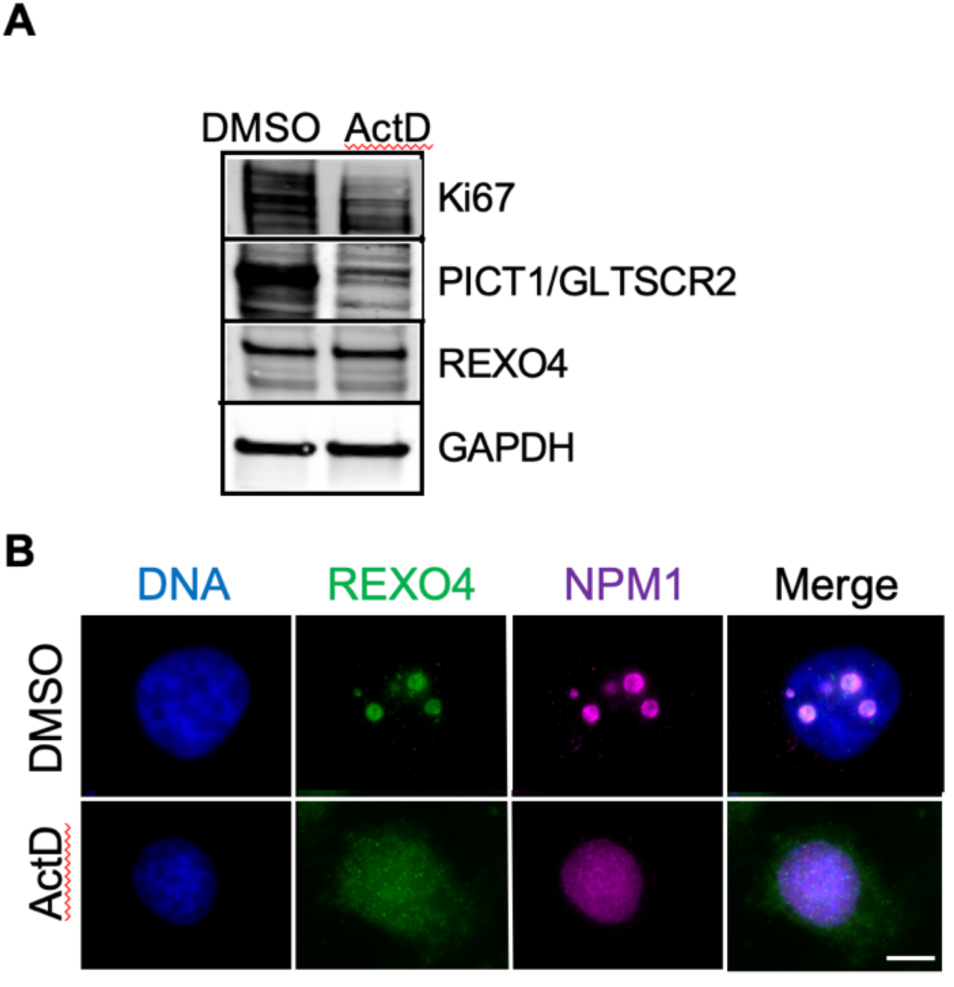
REXO4 is dispersed throughout the cell in response to nucleolar stress. (A) HeLa cells were treated with DMSO control vehicle or 5 nM actinomycin D (ActD) for 24 hours prior to preparation of protein extracts and immunoblot analysis with the indicated antibodies. (B) Immunofluorescence microscopy of HeLa cells treated with control DMSO or 5 nM ActD for 24 hours, fixed, and stained for DNA (Hoechst 33342), REXO4 (anti-REXO4), Ki67 (anti-Ki67), and NPM1 (anti-NPM1). Scale bar indicates 5µm.

## DISCUSSION

Exoribonucleases are critical for ribosome biogenesis due to their ability to process pre-rRNA into mature rRNAs, which are building blocks for ribosome assembly (Dorner *et al*., 2023). Here, we have taken multidisciplinary approaches to characterize the function and regulation of the putative exoribonuclease REXO4. *In silico* analyses showed that REXO4 is conserved among eukaryotes and that multiple isoforms, with varying N-terminal truncations, are likely to be expressed. Further, REXO4 is expressed ubiquitously across diverse tissues and human cell lines and its expression is elevated in cancer. This is consistent with prior reports indicating that REXO4 expression is a biomarker for hepatocellular carcinoma disease progression (Chen *et al*., 2021; Ruan *et al*., 2021). REXO4 subcellular localization studies showed that REXO4 has a dynamic localization pattern during different phases of the cell cycle, localizing to the nucleolus in interphase and re-distributing to the perichromosomal layer of mitotic chromosomes in early mitosis. Further, REXO4’s nucleolar localization was dependent on an N-terminal nucleolar localization sequence, while its localization to the PCL of mitotic chromosomes required the full-length protein and was dependent on Ki67. The REXO4 depletion studies showed that it was important for cellular homeostasis and proliferation, as its depletion led to an arrest in G1/S and a reduction in cell viability.

The localization of REXO4 is consistent with the REXO4 protein-protein association network identified in our mass proteomic studies, as associations with nucleolar-bound proteins, ribosome components, and other proteins involved in rRNA metabolism were enriched. Overall, our data are consistent with a model where REXO4 localizes to the nucleolus and may be important for rRNA processing. This function of REXO4 is likely in turn important for proper ribosome biogenesis, which is critical for cell cycle progression and proliferation (Figure 7).

**Figure 7.**
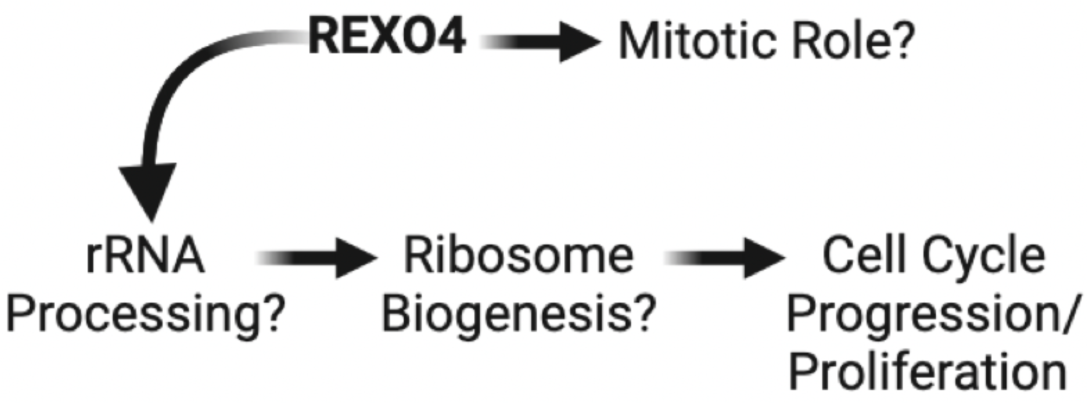
Model for cell cycle regulation by REXO4.

Several of our observations indicate that REXO4 is part of a larger group of nucleolar proteins that share similar properties and behaviors. For example, the long intrinsically disordered region at the N-terminus REXO4 is consistent with other observations that most nucleolar proteins harbor long intrinsically disordered domains (Brangwynne et al, 2011; Nott et al, 2015; Feric et al, 2016). Similarly, the dynamic localization of REXO4 from the nucleolus in interphase to the PCL of mitotic chromosomes is consistent with a previous study that analyzed the subcellular localization of 150 nucleolar proteins during mitosis and showed that about 57% localized to mitotic chromosomes (Stenstrom *et al*., 2020). The function/s of nucleolar proteins that localize to the PCL of mitotic chromosomes is still largely unknown. However, proteins like Ki67 and p120-catenin that have a similar localization pattern to REXO4 have been shown to have roles in regulating mitotic chromosome dynamics and are essential for proper cell division (Cuylen *et al*., 2016; Chow *et al*., 2022). Therefore, it is possible that REXO4 could have a mitotic function that is independent of its role in regulating cell cycle progression. Interestingly, the mislocalization and aggregation of REXO4 to predominantly one end cap of mitotic metaphase chromosomes in Ki67-KO cells is reminiscent of the aggregation of rRNA on metaphase chromosomes that is observed in Ki67-KO cells (Hayashi *et al*., 2017). This was also similar to the localization of nucleolus and PCL-localized proteins Nulceolin and NIFK that cap one end of the metaphase plate in Ki67-depleted cells (Booth *et al*., 2014).

The REXO4 protein harbors a canonical (PX(S/T)P) MAPK/ERK consensus phosphorylation motif within its unstructured N-terminus (13-PSSP-16) and the predicted phosphorylation site Ser15 is phosphorylated in proteome-wide phosphoprotein screens under various settings (Dephoure *et al*., 2008; Olsen *et al*., 2010; Sharma *et al*., 2014). The MAPK/ERK pathway is known to regulate rRNA synthesis (Zhao *et al*., 2003) and multiple aspects of ribonucleic acid metabolism (Bora *et al*., 2021; Ronkina and Gaestel, 2022) and it is possible that REXO4’s activity, stability, localization, or interactions with proteins or RNAs could be modulated by MAPK/ERK and should be explored further. The processing of rRNA is regulated under nucleolar stress conditions (Szaflarski *et al*., 2022) and MAPK/ERK-mediated phosphorylation of REXO4 could be part of this regulation.

## MATERIALS AND METHODS

### Cell culture and drug treatments

HeLa and HEK293T Flp-In T-REx LAP-REXO4 stable cell lines were grown as described previously (Senese *et al*., 2014). For LAP-REXO4 expression, cells were treated with .1 µg/ml doxycycline (Sigma-Aldrich) for 24 hours. For synchronization of cells in early mitosis, cells were treated with 50 µM monastrol (Selleckchem) for 16-hours. For nucleolar damage experiments, cells were treated with 10 nM of actinomycin D (Selleckchem) or matched DMSO control for 24 hours.

### Plasmids, mutagenesis, and generation of stable cell lines

REXO4 full-length (DNASU Clone ID: HsCD000439084), truncation mutants, and point mutants were fused to the C-terminus of GFP using the mammalian expression vector pgLAP1 (Torres *et al*., 2009). REXO4 truncation mutants were generated using primers purchased from IDT. See Supplemental Table S3 for primer sequences. The pgLAP1-REXO4 full-length vector was used to generate a doxycycline inducible HEK293T Flp-In T-REx LAP-REXO4 stable cell line that expresses LAP-REXO4 from a single genomic loci as described in (Torres *et al*., 2009).

### LAP-REXO4 tandem affinity purification and LC-MS/MS analysis

The HEK293T Flp-In T-REx LAP-REXO4 stable cell line was grown in roller bottles and induced with .1μg/ml Dox for 24 hours as described in (Torres *et al*., 2009; Bradley *et al*., 2016). Cells were harvested by agitation and lysed in the presence of protease (Roche), phosphatase (Pierce), de-ubiquitinase (NEM, Enzo Life Sciences), and proteasome inhibitors (MG132, Enzo Life Sciences). LAP-REXO4 purification, eluate resolution by SDS-PAGE, trypsinization of gel slices, and LC-MS/MS analysis at the Harvard Mass Spectrometry and Proteomics Resource Laboratory were as described previously (Vanderwerf *et al*., 2009; Gholkar *et al*., 2016). Identified peptides and their associated data are in Supplemental Table S2.

### Antibodies

The following antibodies were used for immunofluorescence, immunoblotting, and immunoprecipitations: GFP and Ki67 (Abcam), REXO4 and GAPDH (Proteintech), alpha tubulin (Bio-Rad), p62/SQSTM1 (Abcepta), and NPM1 (Abnova). For immunofluorescence microscopy, secondary antibodies conjugated to FITC, CY3, and Cy5 were from Jackson Immuno Research. For immunoblotting, secondary antibodies conjugated were from LI-COR Biosciences. See Supplemental Table S3 for additional information.

### Small interfering RNA (siRNA)-based gene expression knockdown

For siRNA experiments, cells were transfected with 75 nM siRNA against REXO4 or a non-targeting control (Thermo Fisher) using RNAiMAX Transfection Reagent (Invitrogen) as per the manufacturer’s protocol. 72 hours post transfection, cells were collected and whole cell extracts were subjected to immunoblot analysis with the indicated antibodies. See Supplemental Table S3 for additional reagent information.

### Viability assays

Cells were seeded at 5,000 cells/well in black clear-bottom 96-well plates (Corning 3904) and transfected with siRNA 24 hours post-plating. Viability was measured at the indicated time points post-siRNA transfection using the CellTiter-Glo (Promega) assay according to the manufacturer’s protocol and luminescence was measured using a Tecan Infinite M1000 microplate reader as described previously (Xia *et al*., 2019). Experiments were performed in triplicate, averaged, and normalized to negative (non-targeting) siRNA control.

### Flow cytometry

For cell cycle analyses, cells treated with the indicated siRNAs for 72 hours and harvested using TrypLE Express (Thermo Fisher) and washed with PBS. Cold 70% ethanol was added to the cells dropwise, followed by incubation overnight at 4 °C. Fixed cells were centrifuged and stained with 400 μL PI/RNase (Thermo Fisher) solution per million cells. Cells were incubated for 15 minutes at 25 °C. Data acquisition was performed using the Attune NxT Flow Cytometer (Invitrogen). Data was analyzed using FlowJo, gating against smaller cellular debris as well as events consistent with more than one cell per droplet.

### Immunofluorescence microscopy

Immunofluorescence microscopy was carried out essentially as described in (Torres *et al*., 2010). HeLa stable cell lines were induced to express REXO4 or the indicated REXO4 mutants for 24 hours, fixed with 4% paraformaldehyde, permeabilized with 0.2% Triton X-100/PBS, and co-stained with 0.5 μg/ml Hoechst 33342 and indicated antibodies. Images were captured with a Leica Microsystems MICA microscope (63x/1.40 NA oil objective, Leica Microsystems MICA Analysis Package) at room temperature using Hoechst 3342 and secondary FITC, Cy3, and Cy5 antibodies (Jackson Immuno). Images were deconvolved with Leica Microsystems MICA Thunder and Lightning applications using default parameters and exported as TIFF files.

### REXO4 *in silico* analyses

REXO4 ortholog protein sequences were aligned using Clustal Omega with default parameters (Sievers *et al*., 2011). See Supplemental Figure S1 for additional information. Human REXO4 isoform protein sequence alignments were carried out using the UniProt Align module (UniProt, 2023). See Supplemental Figure S2 for additional information. REXO4 gene expression analysis: in diverse non-diseased tissues was performed with the Adult Genotype Tissue Expression (GTEx) Portal (Consortium, 2020) with default parameters; in 1206 human cell lines was conducted with the Human Protein Atlas (Uhlen *et al*., 2015; Jin *et al*., 2023) with default parameters; and in cancer versus matched normal samples was analyzed using the Gene Expression Profiling and Interactive Analysis (GEPIA) web server (Tang *et al*., 2017) with default parameters. See Supplemental Figure S3 for details.

## ACKNOWLEDGMENTS

This material is based upon work supported by the National Science Foundation under Grant Number MCB1912837 and National Institutes of Health under Grant Number GM139539 to J.Z.T.; NIH T32GM145388 fellowships to M.A. and K.J.R.; NIH F31GM154466 fellowship to M.A.; and NIH T32CA009056 grant fellowship to K.M.C. Any opinions, findings, and conclusions or recommendations expressed in this material are those of the authors and do not necessarily reflect the views of the NSF or NIH.

