## Supplementary Material for "Human REXO4 is Required for Cell Cycle Progression"

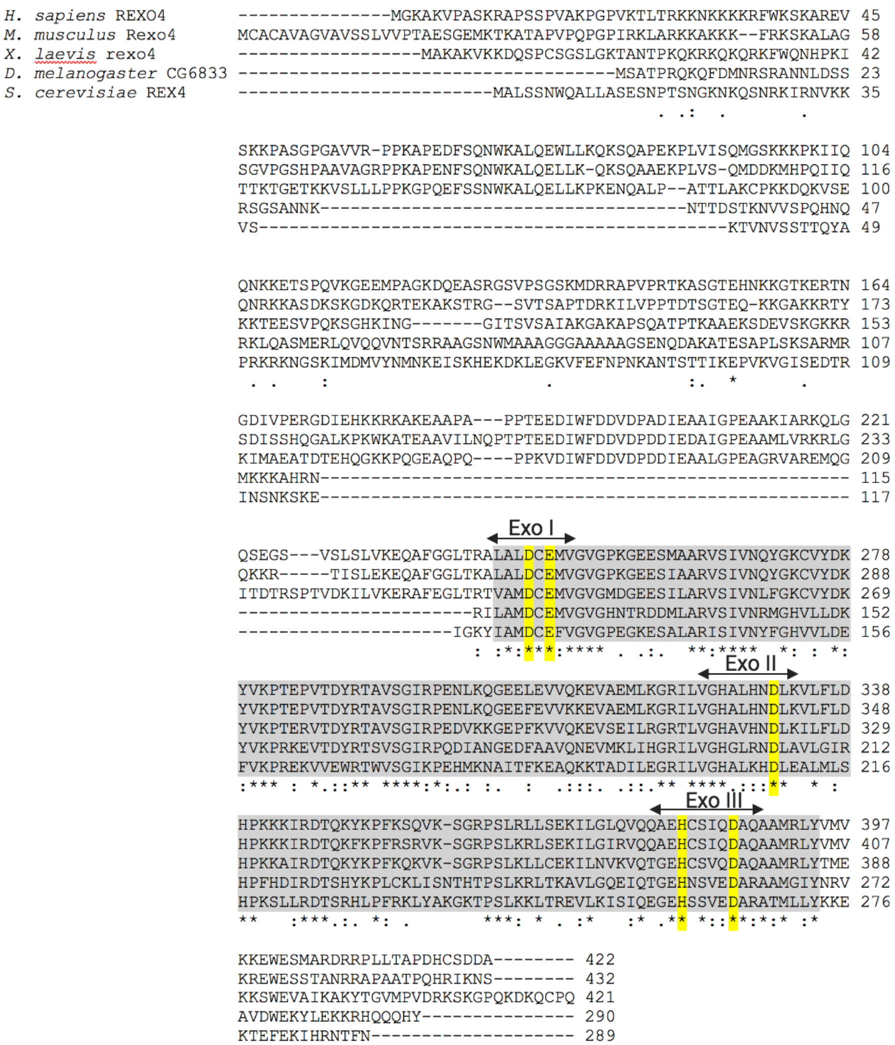


**Figure S1.** REXO4 is conserved among eukaryotes. Amino acid sequence alignment of REXO4 orthologs from common eukaryotic model organisms, *H. sapiens* REXO4: UniProt ID Q9GZR2, *M. musculus* Rexo4: UniProt ID Q6PAQ4, *X. laevis rexo4*: UniProt ID Q91560, *D. melanogaster* CG6833: UniProt ID Q9VUC3, *S. cerevisiae* REX4: UniProt ID Q08237. The number of amino acids is indicated for each protein. The exonuclease domain is highlighted in grey and the three exonuclease motifs (Exo I, Exo II, and Exo III) are marked with arrows. Highlighted in yellow are the conserved DEDDH amino acids characteristic of the DEDDh family of exonucleases. The sequence alignment was conducted in Clustal Omega with default parameters.

```
Isoform 1        MGKAKVPASKRAPSSPVAKPGPVKTLTRKKNKKKKRFWKSKAREVSKKPASGPGAVVRPP 60
Isoform 2        MGKAKVPASKRAPSSPVAKPGPVKTLTRKKNKKKKRFWKSKAREVSKKPASGPGAVVRPP 60
Isoform 3        ------------------------------------------------------------
Isoform 4        ------------------------------------------------------------

Isoform 1        KAPEDFSQNWKALQEWLLKQKSQAPEKPLVISQMGSKKKPKIIQQNKKETSPQVKGEEMP 120
Isoform 2        KAPEDFSQNWKALQEVPRWTGGRQYLAPRPVEQSTIRKEPRKGQ---------------- 104
Isoform 3        ------------------------------------------------------------
Isoform 4        --------------------------------MGSKKKPKIIQQNKKETSPQVKGEEMP 27

Isoform 1        AGKDQEASRGSVPSGSKMDRRAPVPRTKASGTEHNKKGTKERTNGDIVPERGDIEHKKRK 180
Isoform 2        ------------------------------------------------------------
Isoform 3        -----------------MDRRAPVPRTKASGTEHNKKGTKERTNGDIVPERGDIEHKKRK 43
Isoform 4        AGKDQEASRGSVPSGSKMDRRAPVPRTKASGTEHNKKGTKERTNGDIVPERGDIEHKKRK 87

Isoform 1        AKEAAPAPPTEEDIWFDDVDPADIEAAIGPEAAKIARKQLGQSEGSVSLSLVKEQAFGGL 240
Isoform 2        ------------------------------------------------------------
Isoform 3        AKEAAPAPPTEEDIWFDDVDPADIEAAIGPEAAKIARKQLGQSEGSVSLSLVKEQAFGGL 103
Isoform 4        AKEAAPAPPTEEDIWFDDVDPADIEAAIGPEAAKIARKQLGQSEGSVSLSLVKEQAFGGL 147

                    Exo I
Isoform 1        TRALALDCEMVGVGPKGEESMAARVSIVNQYGKCVYDKYVKPTEPVTDYRTAVSGIRPEN 300
Isoform 2        ------------------------------------MVILFQNEGTSSIRSGKLRRQPQP 128
Isoform 3        TRALALDCEMVGVGPKGEESMAARVSIVNQYGKCVYDKYVKPTEPVTDYRTAVSGIRPEN 163
Isoform 4        TRALALDCEMVGVGPKGEESMAARVSIVNQYGKCVYDKYVKPTEPVTDYRTAVSGIRPEN 207

                                           Exo II
Isoform 1        LKQGEELEVVQKEVAEMLKGRILVGHALHNDLKVLFLDHPKKKIRDTQKYKPFKSQVKSG 360
Isoform 2        HPPREELEVVQKEVAEMLKGRILVGHALHNDLKVLFLDHPKKKIRDTQKYKPFKSQVKSG 188
Isoform 3        LKQGEELEVVQKEVAEMLKGRILVGHALHNDLKVLFLDHPKKKIRDTQKYKPFKSQVKSG 223
Isoform 4        LKQGEELEVVQKEVAEMLKGRILVGHALHNDLKVLFLDHPKKKIRDTQKYKPFKSQVKSG 267

                                       Exo III
Isoform 1        RPSLRLLSEKILGLQVQQAEHCSIQDAQAAMRLYVMVKKEWESMARDRRPLLTAPDHCSD 420
Isoform 2        RPSLRLLSEKILGLQVQQAEHCSIQDAQAAMRLYVMVKKEWESMARDRRPLLTAPDHCSD 248
Isoform 3        RPSLRLLSEKILGLQVQQAEHCSIQDAQAAMRLYVMVKKEWESMARDRRPLLTAPDHCSD 283
Isoform 4        RPSLRLLSEKILGLQVQQAEHCSIQDAQAAMRLYVMVKKEWESMARDRRPLLTAPDHCSD 327

Isoform 1        DA 422
Isoform 2        DA 250
Isoform 3        DA 285
Isoform 4        DA 329
```

**Figure S2.** Human REXO4 predicted isoforms. Amino acid sequence alignment of the four predicted human REXO4 isoforms; Isoform 1: NP_065118.2, Isoform 2: NP_001266278.1, Isoform 3: NP_001266279.1, and Isoform 4: NP_001266280.1. Highlighted regions and labeling are the same as in Supplemental Figure S1. Note that Isoform 2 also lacks the Exo I motif of the exonuclease domain. The sequence alignment was performed with the UniProt Align module using default parameters.

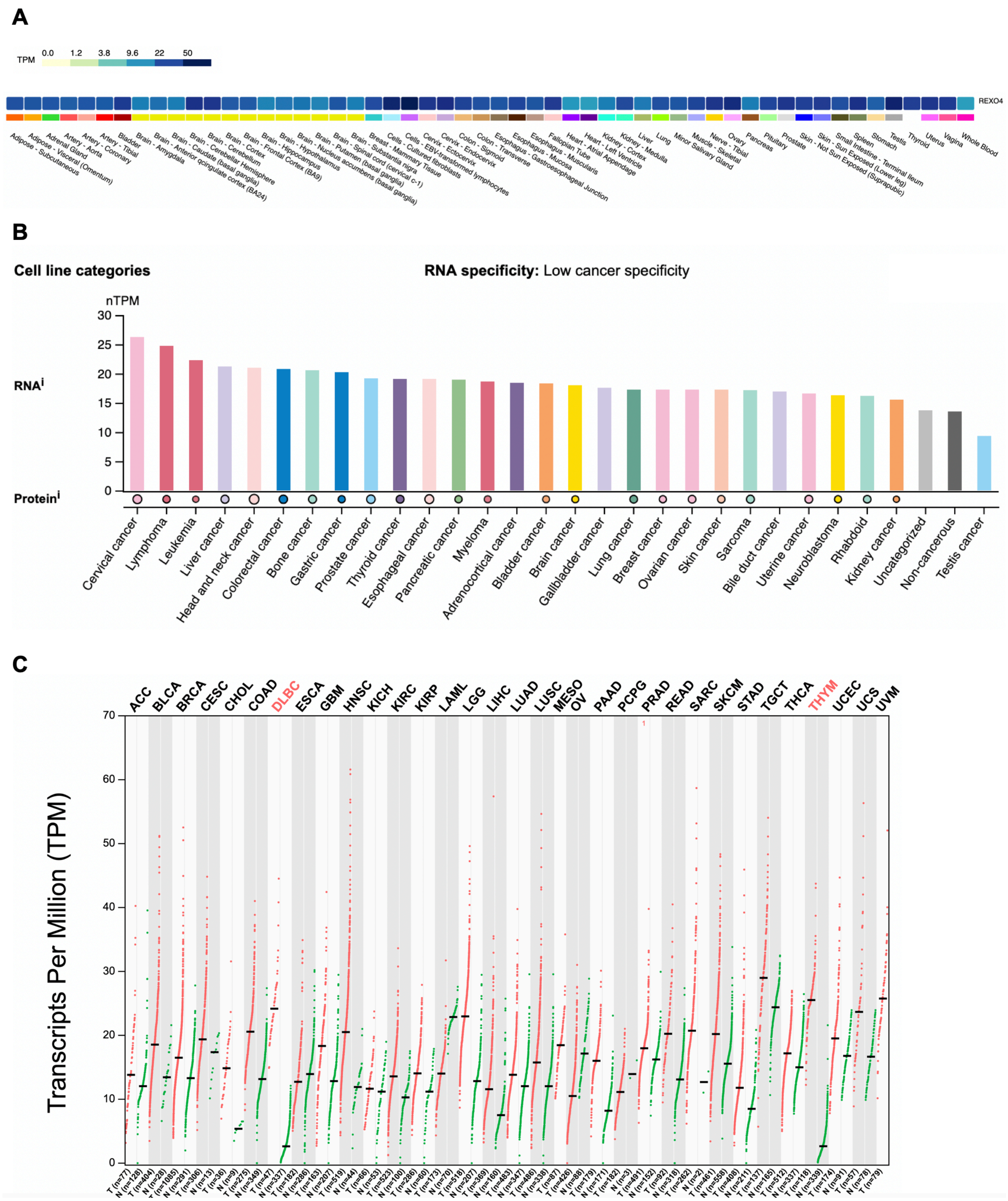


**Figure S3.** REXO4 expression analysis. (A) REXO4 expression analysis across 54 diverse non-diseased tissue types using the Adult Genotype Tissue Expression (GTEx) Portal with default parameters. TPM indicates transcripts per million. (B) Summary of REXO4 expression analysis across 1206 human cell lines, including 1132 cancer cell lines using the Human Protein Atlas with default parameters, nTPM indicates normalized transcript per million. (C) Analysis of REXO4 differential gene expression in various cancers versus matched normal samples with the Gene Expression Profiling and Interactive Analysis (GEPIA) web server using default parameters. See Supplemental Table S1 for a list of cancer types considered and their corresponding abbreviations. The median gene expression levels in transcripts per million (TPM) are on the x-axis and cancer type is on the y-axis.

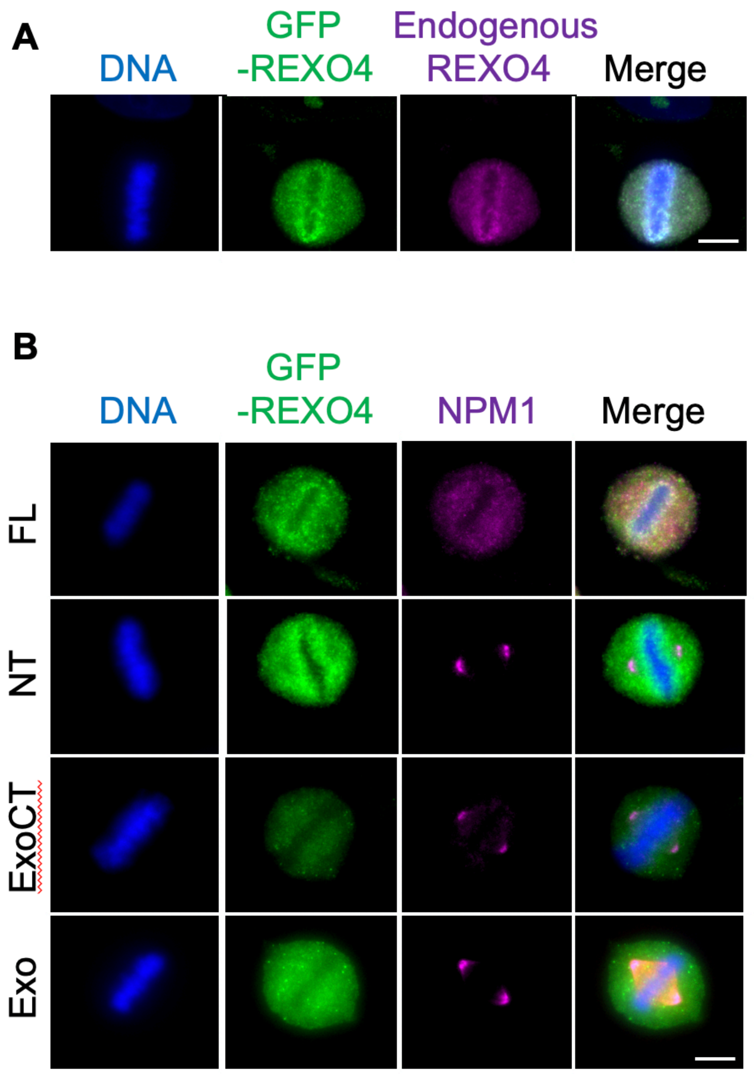


**Figure S4.** Mapping the determinants required for REXO4 localization to metaphase plate chromosomes. (A-B) HeLa cells were transfected with GFP-REXO4-FL, GFP-REXO4-NT, GFP-REXO4-ExoCT, GFP-REXO4-Exo, for 16 hours before fixation. Cells were stained for DNA (Hoechst 33342) and either REXO4 (anti-REXO4), GFP-REXO4 (anti-GFP), or NPM1 (anit-NPM1) as indicated. Bar indicates 5μm.

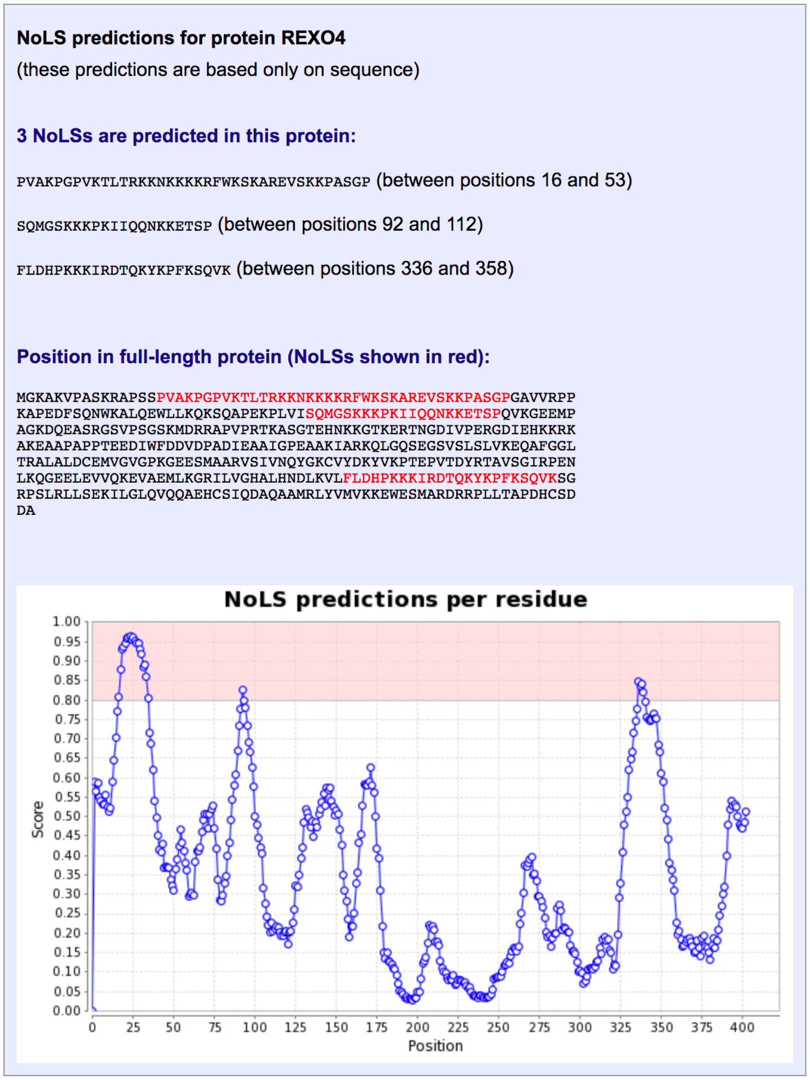


**Figure S5.** REXO4 harbors three predicted nucleolar localization sequences (NoLSs). (A) The human REXO4 Isoform 1 (NP_065118.2) was analyzed with the Nucleolar Localization Sequence Detector (NoD) webserver using default parameters. The amino acid sequences spanning each of the three predicted NoLSs are indicated. The graph shows the NoD NoLS prediction score (y-axis) for each REXO4 amino acid residue (x-axis).

A

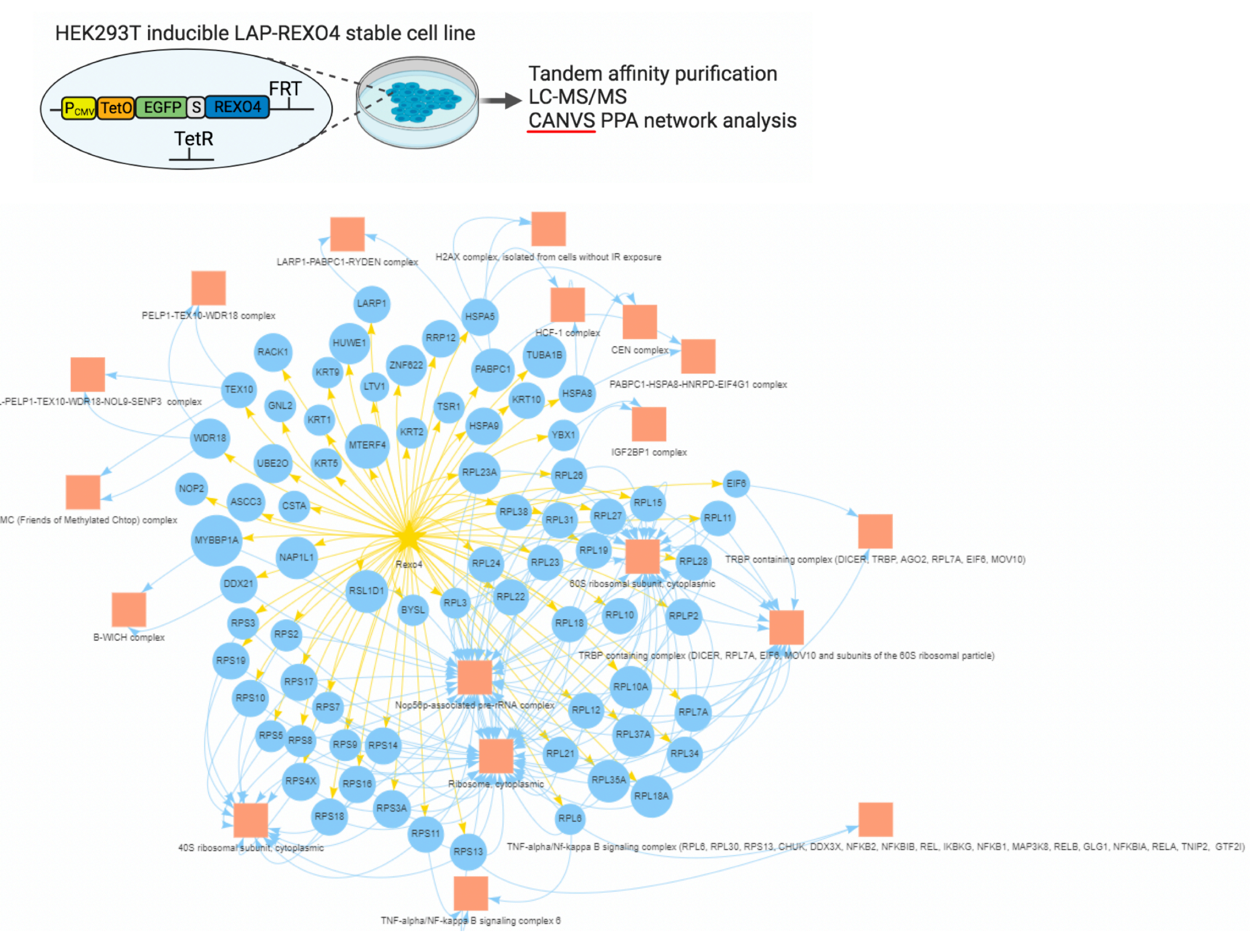


**Figure S6.** REXO4 protein-protein association network. (A) Schematic of HEK293T Flp-In T-REx LAP-REXO4 inducible stable cell line used for REXO4 tandem affinity purification and mass spectrometry analysis to identify associated proteins. (B) Clean Analyze Network Visualization Software (CANVS) analysis of the REXO4 protein-protein interaction network, comprised of proteins identified in the LAP-REXO4 purification analyzed by mass spectrometry. See Supplemental Table S2 for a complete list of REXO4 associating proteins.

**Table S1.** List of cancer types analyzed for REXO4 differential gene expression.

| Abbreviation | Cancer type |
|---|---|
| ACC | Adrenocortical carcinoma |
| BLCA | Bladder Urothelial Carcinoma |
| BRCA | Breast invasive carcinoma |
| CESC | Cervical squamous cell carcinoma & endocervical adenocarcinoma |
| CHOL | Cholangio carcinoma |
| COAD | Colon adenocarcinoma |
| DLBC | Lymphoid Neoplasm Diffuse Large B-cell Lymphoma |
| ESCA | Esophageal carcinoma |
| GBM | Glioblastoma multiforme |
| HNSC | Head and Neck squamous cell carcinoma |
| KICH | Kidney Chromophobe |
| KIRC | Kidney renal clear cell carcinoma |
| KIRP | Kidney renal papillary cell carcinoma |
| LAML | Acute Myeloid Leukemia |
| LGG | Brain Lower Grade Glioma |
| LIHC | Liver hepatocellular carcinoma |
| LUAD | Lung adenocarcinoma |
| LUSC | Lung squamous cell carcinoma |
| MESO | Mesothelioma |
| OV | Ovarian serous cystadenocarcinoma |
| PAAD | Pancreatic adenocarcinoma |
| PCPG | Pheochromocytoma and Paraganglioma |
| PRAD | Prostate adenocarcinoma |
| READ | Rectum adenocarcinoma |
| SARC | Sarcoma |
| SKCM | Skin Cutaneous Melanoma |
| STAD | Stomach adenocarcinoma |
| TGCT | Testicular Germ Cell Tumors |
| THCA | Thyroid carcinoma |
| THYM | Thymoma |
| UCEC | Uterine Corpus Endometrial Carcinoma |
| UCS | Uterine Carcinosarcoma |
| UVM | Uveal Melanoma |

**Table S2.** List of LAP-REXO4 associating proteins identified by mass spectrometry.

| UniProt ID | Description |
|---|---|
| Q7Z6Z7 | HECT, UBA and WWE domain containing E3 ubiquitin protein ligase 1(HUWE1) |
| P52272 | heterogeneous nuclear ribonucleoprotein M(HNRNPM) |
| P11940 | poly(A) binding protein cytoplasmic 1(PABPC1) |
| P08708 | ribosomal protein S17(RPS17) |
| P61353 | ribosomal protein L27(RPL27) |
| P62847 | ribosomal protein S24(RPS24) |
| P68363 | tubulin alpha 1b(TUBA1B) |
| A8K4Z4 | ribosomal protein lateral stalk subunit P0(RPLP0) |
| P46779 | ribosomal protein L28(RPL28) |
| P46778 | ribosomal protein L21(RPL21) |
| P13647 | keratin 5(KRT5) |
| P83881 | ribosomal protein L36a(RPL36A) |
| Q07020 | ribosomal protein L18(RPL18) |
| Q7Z6M4 | mitochondrial transcription termination factor 4(MTERF4) |
| Q92522 | H1.10 linker histone(H1-10) |
| P13645 | keratin 10(KRT10) |
| Q9ULX3 | NIN1 (RPN12) binding protein 1 homolog(NOB1) |
| P04264 | keratin 1(KRT1) |
| P35527 | keratin 9(KRT9) |
| O14730 | RIO kinase 3(RIOK3) |
| P46087 | NOP2 nucleolar protein(NOP2) |
| Q9Y4W2 | LAS1 like ribosome biogenesis factor(LAS1L) |
| Q9HCE1 | Mov10 RNA helicase(MOV10) |
| P55209 | nucleosome assembly protein 1 like 1(NAP1L1) |
| P84098 | ribosomal protein L19(RPL19) |
| P27635 | ribosomal protein L10(RPL10) |
| A8K548 | proline, glutamate and leucine rich protein 1(PELP1) |
| P23396 | ribosomal protein S3(RPS3) |
| P67809 | Y-box binding protein 1(YBX1) |
| P62081 | ribosomal protein S7(RPS7) |
| P38646 | heat shock protein family A (Hsp70) member 9(HSPA9) |
| Q9C0C9 | ubiquitin conjugating enzyme E2 O(UBE2O) |
| P62750 | ribosomal protein L23a(RPL23A) |
| P46783 | ribosomal protein S10(RPS10) |
| P83731 | ribosomal protein L24(RPL24) |
| Q9BQG0 | MYB binding protein 1a(MYBBP1A) |
| P62913 | ribosomal protein L11(RPL11) |
| Q9H4L4 | SUMO specific peptidase 3(SENP3) |
| Q9GZR2 | REX4 homolog, 3'-5' exonuclease(REXO4) |
| P46782 | ribosomal protein S5(RPS5) |

| | |
|---|---|
| P46781 | ribosomal protein S9(RPS9) |
| P16989 | Y-box binding protein 3(YBX3) |
| Q9NQ55 | peter pan homolog(PPAN) |
| P62244 | ribosomal protein S15a(RPS15A) |
| P63173 | ribosomal protein L38(RPL38) |
| P62241 | ribosomal protein S8(RPS8) |
| P61313 | ribosomal protein L15(RPL15) |
| P60866 | ribosomal protein S20(RPS20) |
| Q969S3 | zinc finger protein 622(ZNF622) |
| P62249 | ribosomal protein S16(RPS16) |
| A8K622 | staufen double-stranded RNA binding protein 1(STAU1) |
| Q13895 | bystin like(BYSL) |
| Q9BVP2 | G protein nucleolar 3(GNL3) |
| Q8N726 | cyclin dependent kinase inhibitor 2A(CDKN2A) |
| Q8N3C0 | activating signal cointegrator 1 complex subunit 3(ASCC3) |
| A8K5I0 | heat shock protein family A (Hsp70) member 1A(HSPA1A) |
| A8K5I0 | heat shock protein family A (Hsp70) member 1B(HSPA1B) |
| A2RUM7 | ribosomal protein L5(RPL5) |
| P56537 | eukaryotic translation initiation factor 6(EIF6) |
| P63220 | ribosomal protein S21(RPS21) |
| P62899 | ribosomal protein L31(RPL31) |
| P35908 | keratin 2(KRT2) |
| Q13823 | G protein nucleolar 2(GNL2) |
| P62829 | ribosomal protein L23(RPL23) |
| Q04695 | keratin 17(KRT17) |
| P18077 | ribosomal protein L35a(RPL35A) |
| P15880 | ribosomal protein S2(RPS2) |
| Q9NX58 | Ly1 antibody reactive(LYAR) |
| Q02543 | ribosomal protein L18a(RPL18A) |
| P62263 | ribosomal protein S14(RPS14) |
| P62701 | ribosomal protein S4 X-linked(RPS4X) |
| Q9NXF1 | testis expressed 10(TEX10) |
| P62269 | ribosomal protein S18(RPS18) |
| P62424 | ribosomal protein L7a(RPL7A) |
| Q9Y3U8 | ribosomal protein L36(RPL36) |
| P01040 | cystatin A(CSTA) |
| Q6IPH7 | ribosomal protein L14(RPL14) |
| P62280 | ribosomal protein S11(RPS11) |
| A2A3R6 | ribosomal protein S6(RPS6) |
| Q53GS0 | GTP binding protein 4(GTPBP4) |
| P39019 | ribosomal protein S19(RPS19) |
| P63244 | receptor for activated C kinase 1(RACK1) |
| P62277 | ribosomal protein S13(RPS13) |

| | |
|---|---|
| P27708 | carbamoyl-phosphate synthetase 2, aspartate transcarbamylase, and dihydroorotase(CAD) |
| P30050 | ribosomal protein L12(RPL12) |
| P39023 | ribosomal protein L3(RPL3) |
| P02768 | albumin(ALB) |
| P07437 | tubulin beta class I(TUBB) |
| Q3MIH3 | ubiquitin A-52 residue ribosomal protein fusion product 1(UBA52) |
| Q2NL82 | TSR1 ribosome maturation factor(TSR1) |
| A8K517 | ribosomal protein S23(RPS23) |
| Q9NR30 | DExD-box helicase 21(DDX21) |
| Q96D46 | NMD3 ribosome export adaptor(NMD3) |
| O76021 | ribosomal L1 domain containing 1(RSL1D1) |
| P11142 | heat shock protein family A (Hsp70) member 8(HSPA8) |
| P11021 | heat shock protein family A (Hsp70) member 5(HSPA5) |
| Q9BV38 | WD repeat domain 18(WDR18) |
| P05387 | ribosomal protein lateral stalk subunit P2(RPLP2) |
| P49207 | ribosomal protein L34(RPL34) |
| P61513 | ribosomal protein L37a(RPL37A) |
| A8K3W4 | heterogeneous nuclear ribonucleoprotein U like 1(HNRNPUL1) |
| P32969 | ribosomal protein L9(RPL9) |
| O75683 | surfeit 6(SURF6) |
| P78527 | protein kinase, DNA-activated, catalytic subunit(PRKDC) |
| Q02878 | ribosomal protein L6(RPL6) |
| A8K4C8 | ribosomal protein L13(RPL13) |
| P61247 | ribosomal protein S3A(RPS3A) |
| P02663 | casein alpha-S2(CSN1S2) |
| Q96GA3 | LTV1 ribosome biogenesis factor(LTV1) |
| P62854 | ribosomal protein S26(RPS26) |
| Q6PKG0 | La ribonucleoprotein 1, translational regulator(LARP1) |
| P62906 | ribosomal protein L10a(RPL10A) |
| P35268 | ribosomal protein L22(RPL22) |
| P62861 | FAU ubiquitin like and ribosomal protein S30 fusion(FAU) |
| Q6IB29 | EBNA1 binding protein 2(EBNA1BP2) |
| Q5JTH9 | ribosomal RNA processing 12 homolog(RRP12) |
| O14654 | insulin receptor substrate 4(IRS4) |
| P02533 | keratin 14(KRT14) |

**Table S3.** List of reagents and resources used in this study.

| REAGENT or RESOURCE | SOURCE | IDENTIFIER |
|---|---|---|
| Antibodies | | |
| Rabbit polyclonal anti-REXO4 | Proteintech | Cat# 18890-1-AP; RRID: AB_10643245 |
| Mouse monoclonal anti-NPM1 | Abnova | Cat# H00004869-M01; RRID: AB_10553363 |
| Rabbit monoclonal anti-Ki67 | Abcam | Cat# ab16667; RRID: AB 302459 |
| Rabbit polyclonal anti-PICT1/GLTSCR2 | Proteintech | Cat# 27353-1-AP; RRID: AB_2880852 |
| Chicken polyclonal anti-GFP | Abcam | Cat# ab13970; RRID: AB_300798 |
| Mouse monoclonal anti-GAPDH (clone 1E6D9) | Proteintech | Cat# 60004-1-Ig; RRID: AB_2107436 |
| Rat monoclonal anti-α-Tubulin (clone YOL1/34) | Bio-Rad | Cat# MCA78G; RRID: AB_325005 |
| Donkey polyclonal anti-Rat IgG (H+L), Cy3 AffiniPure | Jackson ImmunoResearch Labs | Cat# 712-165-153; RRID: AB_2340667 |
| Donkey polyclonal anti-Chicken IgY (IgG) (H+L), Fluorescein (FITC) AffiniPure | Jackson ImmunoResearch Labs | Cat# 703-095-155; RRID: AB_2340356 |
| Donkey polyclonal anti-Rabbit IgG (H+L), Fluorescein (FITC) AffiniPure | Jackson ImmunoResearch Labs | Cat# 711-095-152 RRID: AB_2315776 |
| Donkey polyclonal anti-Mouse IgG (H+L), Cy5 AffiniPure | Jackson ImmunoResearch Labs | Cat# 715-175-150 RRID: AB_2340819 |
| Donkey polyclonal anti-Rabbit IgG (H+L), Cy5 AffiniPure | Jackson ImmunoResearch Labs | Cat# 711-175-152 RRID: AB_2340607 |
| Donkey polyclonal anti-Mouse IgG (H+L), IRDye 800CW | LI-COR Biosciences | Cat# 926-32212; RRID: AB_621847 |
| Donkey polyclonal anti-Rabbit IgG (H+L), IRDye 680RD | LI-COR Biosciences | Cat# 926-68073; RRID: AB_10954442 |
| Donkey polyclonal anti-Chicken IgG (H+L), IRDye 680RD | LI-COR Biosciences | Cat# 925-68075; RRID: AB_2814924 |
| Chemicals, Peptides, and Recombinant Proteins | | |
| Monastrol | Selleckchem | Cat# S8439; CAS: 329689-23-8 |
| MI-181 | Enamine | Cat# Z46083298 CAS: N/A |
| Actinomycin-D, ActD | Selleckchem | Cat# S8964; CAS: 50-76-0 |
| Hygromycin B | Thermo Fisher Scientific | Cat# 10687010 |
| Doxycycline | Sigma-Aldrich | Cat# D9891; CAS: 24390-14-5 |
| MG132 | Millipore Sigma | Cat# 474790; CAS: 133407-82-6 |
| Halt Protease Inhibitor Cocktail | Thermo Fisher Scientific | Cat# 87786 |
| Phosphatase Inhibitor Cocktail 2 | Sigma-Aldrich | P5726 |

| | | |
|---|---|---|
| Phosphatase Inhibitor Cocktail 3 | Sigma-Aldrich | P0044 |
| Hoechst 33342 | Thermo Fisher Scientific | Cat# H1399; CAS: 23491-52-3 |
| ProLong Gold Antifade Mountant | Thermo Fisher Scientific | Cat# P36934 |
| Lipofectamine RNAiMAX | Thermo Fisher Scientific | Cat# 13778150 |
| FuGENE 6 Transfection Reagent | Promega | Cat# E2691 |
| FuGENE HD Transfection Reagent | Promega | Cat# E2311 |
| S-protein Agarose | Millipore Sigma | Cat# 69704-4 |
| Critical Commercial Assays | | |
| Gateway LR Clonase II Enzyme mix | Thermo Fisher Scientific | Cat# 11791020 |
| Gateway BP Clonase II Enzyme mix | Thermo Fisher Scientific | Cat# 11789020 |
| QIAprep Spin Miniprep Kit | QIAGEN | Cat# 27106 |
| Monarch Total RNA Miniprep Kit | New England Biolabs | Cat# T2010S |
| FxCycle PI/RNase Staining Solution | Thermo Fisher Scientific | Cat# F10797 |
| Experimental Models: Cell Lines | | |
| HeLa | ATCC | Cat# CCL-2; RRID: CVCL_0030 |
| HeLa Ki67 knock out | Gerlich Lab | (Cuylen *et al.*, 2016) |
| Inducible HEK293T Flp-In T-REx LAP-REXO4-FL | This paper | N/A |
| Inducible HeLa Flp-In T-REx | Taylor Lab | (Tighe *et al.*, 2008) |
| Inducible HeLa Flp-In T-Rex- REXO4-FL | This paper | N/A |
| Inducible HeLa Flp-In T-Rex- REXO4-NT | This paper | N/A |
| Inducible HeLa Flp-In T-Rex- REXO4-ExoCT | This paper | N/A |
| Inducible HeLa Flp-In T-Rex- REXO4-Exo | This paper | N/A |
| Inducible HeLa Flp-In T-Rex- REXO4-NoLS1 | This paper | N/A |
| Inducible HeLa Flp-In T-Rex- REXO4-NoLS1+2 | This paper | N/A |
| Oligonucleotides | | |
| siRNA 1 targeting REXO4 | Thermo Fisher Scientific | Cat# 4392420; siRNA ID: s224450 |
| siRNA 2 targeting REXO4 | Thermo Fisher Scientific | Cat#: 4392420; siRNA ID: s32694 |
| Primer for REXO4-NT: Fwd 5'-GGGGACAAGTTTGTACAAAAAAGCAGGCTTCGAAGGAGATAGAACCATGGGGAAGGCGAAGGTCCCCGC-3' | Eurofins Genomics | N/A |
| Primer for REXO4-NT: Rev 5'-GGGGACCACTTTGTACAAGAAAGCTGGGTCCTATCTTGTCAGGCCGCCGAA-3' | Eurofins Genomics | N/A |
| Primer for REXO4-ExoCT: Fwd 5'-GGGGACAAGTTTGTACAAAAAAGCAGGCTTCGAAGGAGATAGAACCATGGCCTTAGCCTTGGACTGTGA-3' | Eurofins Genomics | N/A |

| | | |
|---|---|---|
| Primer for REXO4-ExoCT: Rev 5'-AGTCGAGGCTGATCAGCGGGTTTAAACGGGCCCTCTAGACTCGACTAGGCGTCGTCACTGCAGT-3' | Eurofins Genomics | N/A |
| Primer for REXO4-Exo: Fwd 5'-GGGGACAAGTTTGTACAAAAAAGCAGGCTTCGAAGGAGATAGAACCATGGCCTTAGCCTTGGACTGTGA-3' | Eurofins Genomics | N/A |
| Primer for REXO4-Exo: Rev 5'-GGGGACCACTTTGTACAAGAAAGCTGGGTCCTAGTACAGCCTCATTGCTGCCT-3' | Eurofins Genomics | N/A |
| Primer for REXO4-NoLS 1+2: Fwd 5'-ACAAGTTTGTACAAAAAAGCAGGCTTCATGCCCGTGGCTAAGCCGGGTCCT | Eurofins Genomics | N/A |
| Primer for REXO4-NoLS 1+2: Rev 5'-ACCACTTTGTACAAGAAAGCTGGGTCTAGGGGCCGCTTGCTGGCTTCTT | Eurofins Genomics | N/A |
| Primer for REXO4-NoLS 1: Rev 5'-ACCACTTTGTACAAGAAAGCTGGGTCTAAGGCGAGGTCTCTTTTTTGTT | Eurofins Genomics | N/A |
| Recombinant DNA | | |
| pDONR221-REXO4 | DNASU Plasmid Repository | Clone ID: HsCD000439084 |
| pgLAP1-REXO4 | This paper | N/A |
| pgLAP1-REXO4-NT | This paper | N/A |
| pgLAP1-REXO4-ExoCT | This paper | N/A |
| pgLAP1-REXO4-Exo | This paper | N/A |
| pgLAP1-REXO4-NoLS1 | This paper | N/A |
| pgLAP1-REXO4-NoLS1+2 | This paper | N/A |
| Software and Algorithms | | |
| GraphPad Prism 10 | GraphPad | RRID: SCR_002798 |
| BioRender | BioRender | RRID: SCR_018361 |
| Clean Analyze Network Visualization Software (CANVS) | Torres Lab | (Velasquez *et al.*, 2021) |
| Nucleolar Localization Sequence Detector (NoD) | Barton Lab | (Scott *et al.*, 2011) |
| Clustal Omega | Higgins Lab | (Sievers *et al.*, 2011) |
| Uniprot Align | N/A | (UniProt, 2023) |
| FlowJo | BD Biosciences | RRID: SCR_008520 |
